# Event timing precision in stochastic gene expression with bursty production

**DOI:** 10.64898/2026.10.25.761601

**Authors:** Sayeh Rezaee, Pavol Bokes, Abhyudai Singh

## Abstract

The timing of many cellular events is controlled by the accumulation of a regulatory molecule to a threshold level. This timing can be characterized by the first-passage time (FPT), defined as the random time required to reach the threshold for the first time. In stochastic gene expression, fluctuations in molecular production and degradation lead to variability in event timing. Feedback regulation of the production rate is a natural mechanism for reducing FPT variability. Here, we first investigate how degradation reshapes the optimal feedback law for minimizing timing variability across low- and high-threshold regimes. Using backward equations and Lagrange multiplier formulation, we derive the corresponding optimality conditions under a fixed mean FPT constraint. Exact results for the low-threshold case and asymptotic analysis in the high-threshold, small-degradation regime show that the optimal production rates increase approximately linearly with molecule number before decreasing near the threshold. For larger degradation rates, the optimality conditions develop multiple solution branches and fold behavior, and the optimal production rate structure can become nonmonotonic. We then extend the framework to bursty production and numerically characterize the optimal feedback in both low- and high-threshold regimes. The results show how degradation and bursty synthesis qualitatively reshape the structure of the optimal feedback for stochastic cellular event timing.

## I. Introduction

In cellular systems, the timing of many regulatory processes is determined by the time at which the level of a signaling molecule reaches a critical threshold and triggers a downstream event [1]–[6]. This mechanism is involved in a variety of biological processes, including cell cycle progression and cell division, where threshold levels of cyclin-dependent kinase activity regulate commitment and phase transitions [7]–[10]. Another well known example is bacteriophage *λ*, in which holin proteins accumulate in the bacterial inner membrane until their abundance triggers membrane permeabilization and host lysis [11]–[13].

Molecular interactions are inherently stochastic, and many regulatory species are present at low copy numbers. As a result, the time required to reach the threshold varies from cell to cell [14]–[17]. This intrinsic noise introduces variability into processes that depend on precise molecular accumulation. Consequently, understanding such systems requires not only determining the mean threshold-crossing time but also quantifying the cell-to-cell timing variability caused by stochastic fluctuations.

In this work, we analyze a stochastic gene expression model in which protein molecules are produced and degraded randomly, and an event is triggered when the protein level first reaches a target threshold. The first-passage time (FPT) framework [18]–[21] provides a natural description of this problem, with event timing defined as the random time required for a regulatory molecule to reach a prescribed threshold [22]–[27]. In this framework, the mean FPT determines the average event time, while higher-order moments quantify timing precision across cells. We then study how this precision can be optimized while preserving a desired mean FPT. To address this question, we consider a biologically plausible mechanism: self-regulation of protein synthesis, in which the production rate depends on the current protein level [28]–[32], and examine the feedback strategy that minimizes fluctuations in event timing.

Prior studies have established important results on the control of gene expression timing. In the absence of degradation, the optimal strategy for minimizing timing noise under a fixed mean FPT is constant production, showing that feedback is not always beneficial [33]. This framework was later extended to continuous dilution, where increasing feedback can partially restore timing precision [34]. More recently, stochastic degradation has been studied under restricted feedback classes, showing that positive feedback is optimal within linear regulation, while a mixed feedback profile can further reduce timing fluctuations when these restrictions are relaxed [35]. Beyond direct feedback optimization, related studies have examined how regulatory mechanisms influence timing precision, including regulation by dynamic activators and repressors, gene-regulatory architecture and network size, modulation of burst frequency or burst size by external signals [36]–[40] and multi-gene cascades [41]–[44]. FPT approaches have also been used to characterize time-dependent transcription and post-transcriptional regulation by small RNAs [45]–[48], as well as bacteriophage lysis timing [49]–[51].

Despite the broad literature on this topic, the structure of the optimal state-dependent production rate in the presence of stochastic degradation remains incompletely characterized, particularly beyond prescribed feedback forms and under bursty synthesis. In this work, we optimize the production rate at each molecular state under a fixed mean FPT constraint. We derive analytical results for the nonbursty model in low-threshold and weak-degradation regimes and complement them with numerical solutions at larger degradation, before extending the framework to bursty production to determine how burst statistics reshape the optimal feedback strategy.

## II. Modeling Event Timing as a First-Passage Time Problem

We aim to model the timing of a cellular event driven by stochastic gene expression. In this model, the event is triggered when the protein number first reaches a prescribed threshold. Let *x*(*t*) ∈ {0, 1, 2, … *}*denote the copy number of the protein at time *t*. The dynamics of *x*(*t*) is modeled as a birth-death process with state-dependent synthesis. Hence, when *x*(*t*) = *i*, the probability of producing one molecule during an infinitesimal interval (*t, t* + *dt*] is given by

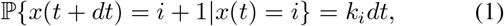

where *k*_*i*_ is the production rate at state *i*. Consequently, the time required to transition from *i* to *i* + 1 molecules is exponentially distributed with mean 1*/k*_*i*_. In this context, a decreasing dependence of *k*_*i*_ on *i* represents negative feedback, whereas an increasing dependence corresponds to positive feedback. More general nonmonotonic forms of *k*_*i*_ capture more complex feedback strategies. Note the special case *k*_*i*_ = *k* corresponds to synthesis without feedback.

Degradation is assumed to occur independently for each molecule, such that

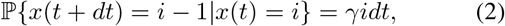

where *γ* is the degradation rate. Let *X* denote a positive integer threshold in molecular copy number that triggers the event of interest. The first-passage time is then identified as

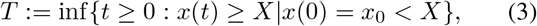

which corresponds to the first time the random process *x*(*t*) reaches *X* when starting from an initial condition *x*_0_ at time *t*=0 (see Fig. 1).

**Fig. 1:**
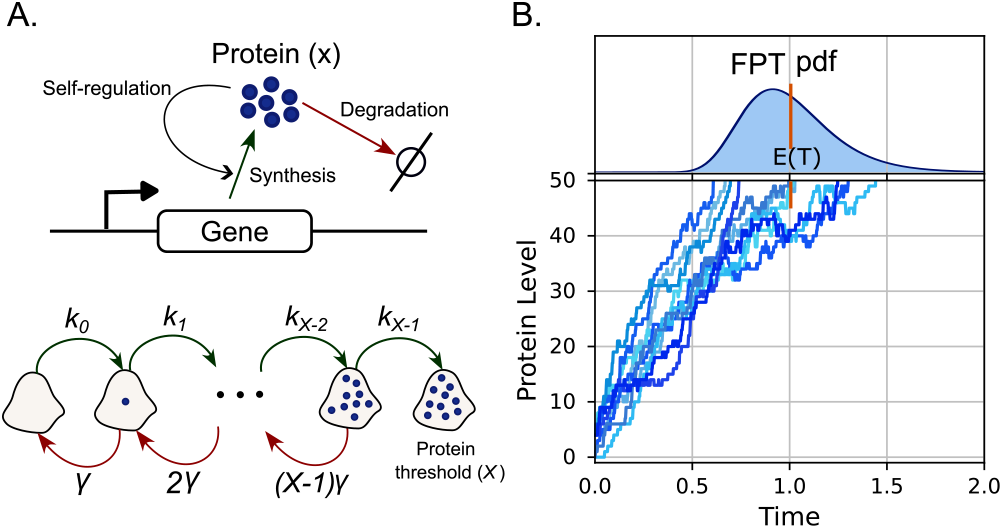
Stochastic timing of threshold-triggered events in a feedback-regulated gene expression model. **A**. Schematic of the gene expression model. Protein molecules are produced and degraded stochastically, with self-regulation implemented through a state-dependent synthesis rate *k*_*i*_, depending on current protein copy number *i*. Protein degradation rate is *γ*. The cellular event is triggered when protein count first reaches the absorbing threshold state *X*. In the shown example, the threshold is *X* = 10. **B**. Example protein copy-number trajectories showing variability in threshold-crossing time for a 50-molecule threshold. The corresponding FPT probability density is shown above, with mean E(*T*) = 1.

For a given *X* and degradation rate *γ*, the aim is to identify the optimal feedback law *k*_*i*_, *i* ∈{0, 1, …, *X −* 1} for the production that minimizes fluctuations in *T* subject to a fixed mean first-passage time, denoted by *C*. Precisely, for given *γ ≥* 0, *x*(0) = 0, and *C >* 0, we seek 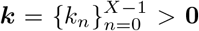 such that

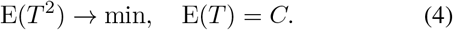

Here E(.) denotes expectation of a random variable. Equivalent objective functions include the variance of the FPT, i.e., E(*T* ^2^) − E^2^(*T*) *→*min with E(*T*) = *C*, or the squared coefficient of variation:

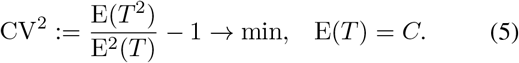

The constrained optimization problem (4) leads to the Lagrangian

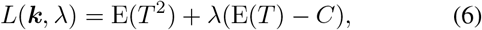

where *λ* is the Lagrange multiplier associated with the fixed mean FPT constraint. The corresponding Lagrange equations are given by

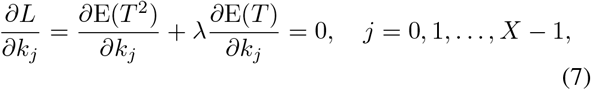

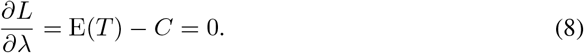

This system of *X* + 1 equations determines *X* + 1 unknowns: candidate values of the *X* production rates and the Lagrange multiplier. The minimizing solution of the constrained problem is denoted by the optimal rate vector 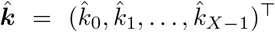 with the corresponding multiplier 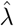.

## III. Optimal Feedback Strategy in Non-bursty Production Model

To solve (7)–(8), we must evaluate the derivatives of the Lagrangian for arbitrary parameter values. Although (7)–(8) are formulated for the FPT starting from the initial condition *n* = 0, this quantity is coupled to FPTs starting from other initial states. Let

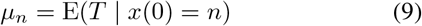

denote the mean FPT starting from state *n*. The vector 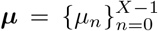 satisfies the *backward equation* [52], [53] (Appendix A)

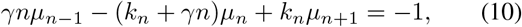

for *n* ∈ {0, 1, …, *X −*1}, with the boundary condition *µ*_*X*_ = 0 at the threshold state. The backward equation (10) forms a linear tridiagonal system of *X* equations in the *X* unknowns ***µ***. It can be solved in *O*(*X*) operations, using the Thomas algorithm, a specialized form of Gaussian elimination for tridiagonal systems [54]. In our implementation, we employ the banded matrix solver scipy.linalg.solve banded from the scientific library SciPy in the Python programming language, which also has *O*(*X*) computational complexity. After computing the solution, we substitute its zeroth component *µ*_0_ = E(*T*) into (8) to evaluate the multiplier derivative of the Lagrangian.

The derivative of the Lagrangian with respect to the production rate is expressed in (7) in terms of derivatives of the FPT moments. To derive equations for these derivatives, we differentiate the backward equation (10) with respect to *k*_*j*_, *j* = 0, 1, …, *X −* 1, which yields the *derivative backward equations*

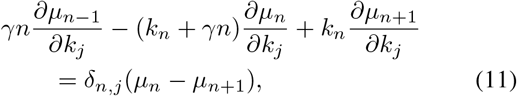

for *n* ∈ {0, 1, …, *X −*1}. Here, *δ*_*n,j*_ denotes the Kronecker delta, i.e., *δ*_*n,j*_ = 1 if *n* = *j* and *δ*_*n,j*_ = 0 otherwise. At the threshold state, we have 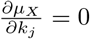.

For a fixed *j* ∈ {0, 1, …, *X* −1}, equation (11) again forms a linear tridiagonal system of *X* equations in the *X* unknowns 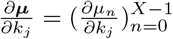. Note that the solution ***µ*** of (10) appears on the right-hand side of (11).

Using this approach, we can solve the constrained optimization problem in the two regimes considered here. We first analyze the threshold *X* = 3, which is the simplest case and allows explicit analytical characterization of the optimal feedback structure. We then study the large-threshold regime, where asymptotic methods yield analytical insight into the form of the optimal solution for larger molecule numbers.

### A. Three-Molecule Threshold

To study the simplest form of the optimization problem, we consider the threshold *X* = 3. Note *X* = 2 can be solved explicitly within the same framework and yields identical optimal rates, reflecting its relatively simple structure [35]. On the other hand, *X* = 3 allows more general feedback profiles to arise. The corresponding Lagrange equations for this case are obtained using backward equations as

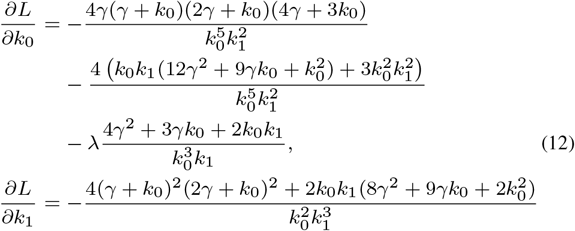

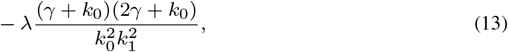

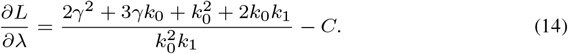

To make the optimization problem analytically tractable, we impose the relation *k*_0_ = *k*_2_, motivated by the approximately symmetric pattern observed in numerical solutions of the fixed-mean-FPT optimization problem when no additional constraints are imposed on the production rates, as reported in our previous work [35]. Under this assumption, the Lagrange optimality conditions can be reduced to a single equation for the remaining rate parameter

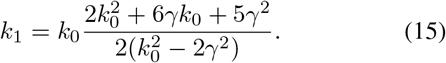

The optimum is then the unique real root of the following polynomial satisfying *k*_0_*C >* 2, which minimizes *E*(*T* ^2^) under the constraint *E*(*T*) = *C*.

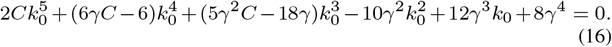

The resulting equation is quintic and therefore does not admit a general closed-form solution, so the optimal value of *k*_0_ is determined numerically. However, in the small-degradation regime, *γC ≪*1, an analytical approximation can be obtained by expressing the solution as a power-series expansion in *γC*.

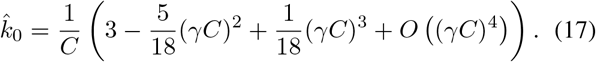

Here, *O* (*γC*)^4^ denotes terms of fourth order and higher in the small parameter *γC*.

Fig. 2 illustrates the optimal production rates for the three-molecule threshold case. The numerical solution of (16) shows that the optimal intermediate rate 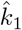 is consistently larger than both 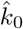 and 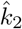. The optimal feedback profile depends strongly on the degradation rate *γ*:

**Fig. 2:**
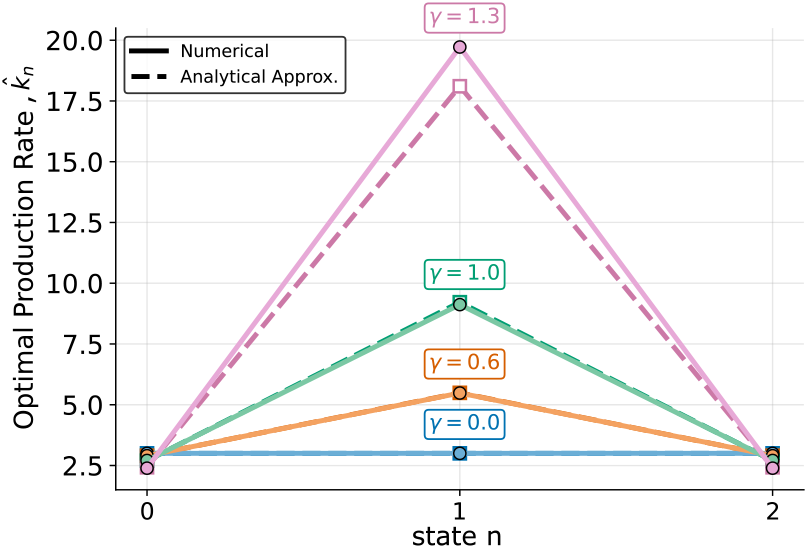
Optimal production rates for a threshold of three molecules in nonbursty gene expression model. Optimal rates are obtained subject to the fixed mean FPT, *C* = 1 with the additional constraint 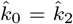. This relation is motivated by the approximate equality of these rates observed numerically when the production rates are optimized without additional constraints. Rates are shown for degradation rates *γ* ∈{0, 0.6, 1, 1.3}. Solid lines represent the numerical optimum obtained in Section III, while dashed lines denote the corresponding analytical approximation valid for *γC ≪* 1 shown in (17).

As *γ* increases, 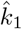 also rises, indicating that degradation increasingly shapes the structure of the optimal feedback law. For sufficiently strong degradation, the noise-minimizing strategy shifts production toward the intermediate state, thereby compensating for degradation-induced backward fluctuations. In the regime *γC≪* 1, the analytical approximation shown in (17) closely matches the numerical solution, confirming the validity of the asymptotic result.

### B. Large-Molecule Threshold

To compute the mean FPT and its derivatives for this case, using (10) and (11) we must solve *X* + 1 tridiagonal systems, each requiring *O*(*X*) operations, resulting in an overall computational complexity of *O*(*X*^2^). An analogous procedure applies to the second moment, with the same *O*(*X*^2^) complexity. The backward-equation framework thus renders the evaluation of Lagrange derivatives (7)–(8) computationally feasible for moderately large values of *X*.

For small values of *γ >* 0, the Lagrange equations can be solved using regular perturbation theory (Appendix B)

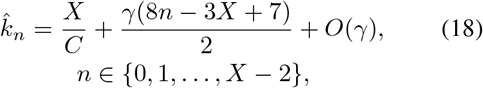

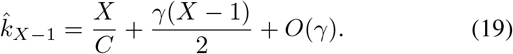

For *γ* = 0 (no degradation), (18)–(19) reduce to the constant solution reported in [35]. The perturbation terms in (18)–(19) introduce a linear profile in *n* with a pronounced drop in the final component (Fig. 3).

**Fig. 3:**
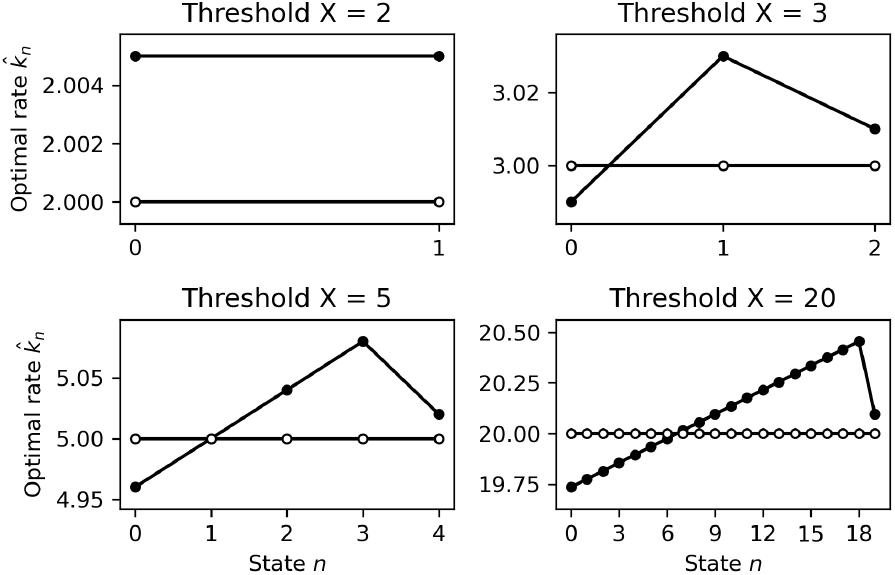
Optimal production rates for zero and small degradation in nonbursty gene expression model. The rates are expressed in equations (18)–(19) and are plotted for different values of threshold *X* −{2, 3, 5, 20}. White circles correspond to no degradation (*γ* = 0), while black circles correspond to slow degradation (*γ* = 0.01). Horizontal axis shows the state index *n*, corresponding to 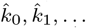 and vertical axis shows the value of associated optimal production rate 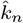. Mean FPT is fixed to *C* = 1.

For larger values of *γ >* 0, solutions of the Lagrange equations are obtained by numerical continuation [55]. This produces a solution branch 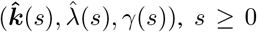, such that 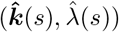 satisfies the Lagrange equations (7)–(8) for *γ* = *γ*(*s*).

The branch originates from the no-degradation solution with *γ*(0) = 0 and 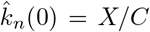. For small *γ*, the linear regime (18)–(19) applies. For *X≥* 3, the optimal rate 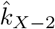, corresponding to two states below the threshold, begins to increase rapidly as *γ* grows further. For *X* = 3, there exists a finite value *γ* = *γ*_blowup_ = *γ*(*s*_blowup_) such that

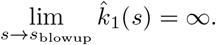

Numerical evidence suggests that 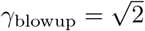 For *X ≥*4, there exists a fold point *γ* = *γ*_fold_ = *γ*(*s*_fold_) satisfying *γ*^*′*^(*s*_fold_) = 0. For *γ < γ*_fold_, two distinct solutions of the Lagrange equations coexist and merge at the fold point.

We expect that the fold point changes the character of the solution, i.e., the solution with the smaller value of 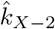 is a minimizer (Fig. 4, solid lines), whereas the solution with the larger value of 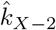 is a saddle point (Fig. 4, dashed lines). The optimal solution oscillates for large *γ* (Fig. 5).

**Fig. 4:**
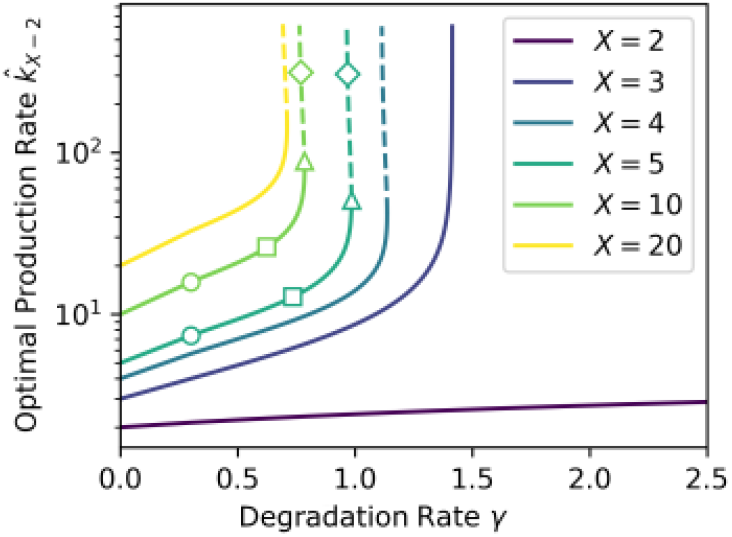
Parametric continuation of the optimal solution for large degradation rates *γ >* 0 in the non-bursty gene expression model. The optimal rate 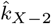 is shown as a function of degradation rate *γ* for thresholds *X* ∈ {2, 3, 4, 5, 10, 20}, with mean FPT fixed at *C* = 1. For *X >* 2, 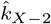 corresponds to the peak optimal production rate; for *X* = 2, the two optimal rates are equal. Optimal production rates are obtained by numerically solving the Lagrange derivatives (7)–(8).

**Fig. 5:**
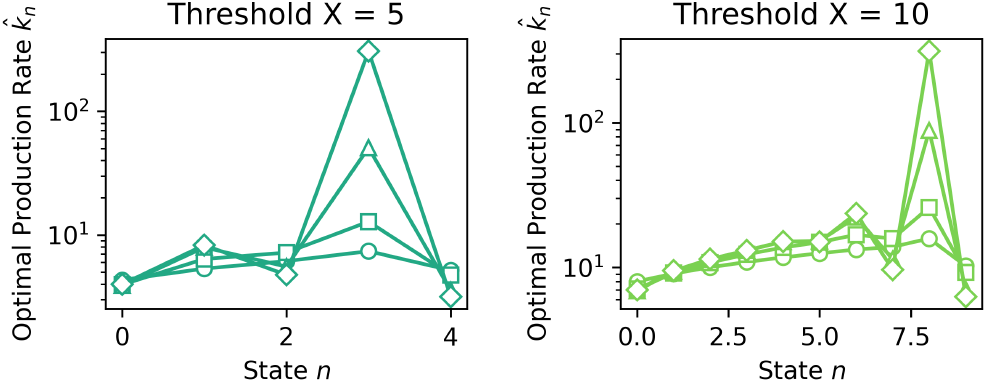
Parametric continuation of the optimal solution for thresholds *X* = 5 and *X* = 10 in the non-bursty gene expression model. Values of *γ* are highlighted by the corresponding markers in Fig. 4. The horizontal axis shows the state index *n*, corresponding to 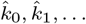 and the vertical axis shows the value of the associated optimal production rate 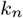 obtained by numerically solving the Lagrange derivatives (7)–(8). The mean FPT is fixed to *C* = 1.

## IV. Optimal Feedback Strategy in Bursty Production Model

Since bursty gene expression is widely observed across biological systems [56]–[58], it is important to determine how burst-driven synthesis fluctuations reshape the optimal strategy for controlling event timing. To incorporate bursty synthesis, the birth process is modeled such that each synthesis event produces a random number of molecules. When the system is in state *i*, burst events occur at a rate *k*_*i*_ such that

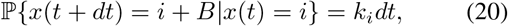

where *B* is a geometrically distributed random variable representing the protein burst size. The probability mass function of *B* is given by

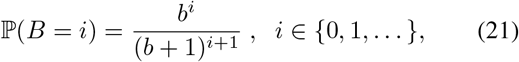

with *b >* 0 denoting the mean production burst size. production burst size. Degradation is unchanged and occurs through single-molecule loss. Note that the nonbursty model corresponds to deterministic unit-size production events, i.e., *B* = 1 with probability one.

We formulate the optimization problem for the case *X* = 3 analogously to the nonbursty model in Section III, using backward equations and Lagrange optimality conditions modified to account for burst transitions. Fig. 6A shows the numerical results for optimal production rates, together with the corresponding nonbursty optimum under the same fixed-mean-FPT constraint, with no additional constraints. Unlike the nonbursty case, bursty optimum does not satisfy 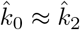, showing that burst production qualitatively changes the optimal feedback structure. The most pronounced difference is in 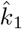 it is higher in the bursty model at low degradation, but this ordering reverses at higher degradation *γ* = 1.3. The nonbursty model requires a substantially larger *k*_1_ to compensate for molecule loss through single-molecule production events, whereas bursty synthesis can recover lost progress through larger production jumps, achieving the same mean FPT with lower intermediate production rates. To examine the effect of burst size on the optimal production rate structure for *X* = 3, Fig. 6B shows that increasing the mean burst size *b* lowers the optimal production rates, since larger bursts move the system closer to the threshold in each synthesis event, reducing the production rates required to maintain the prescribed mean FPT.

**Fig. 6:**
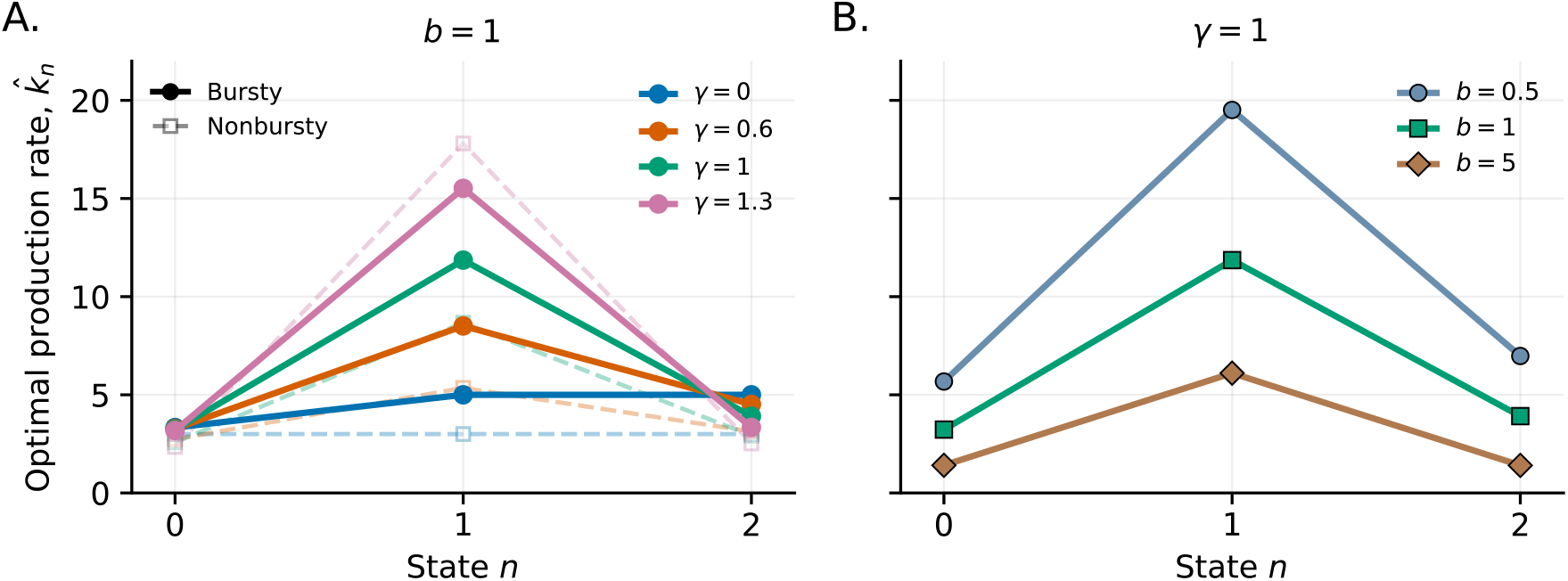
Degradation and burst size reshape the optimal production rates for a three-molecule threshold in bursty gene expression model. **A**. Optimal production rates for the bursty model with mean burst size *b* = 1 and degradation rates *γ* ∈ {0, 0.6, 1, 1.3}. The optimal structure is nonmonotonic, with 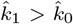 followed by a decrease in 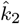. Dashed lines show the corresponding numerical optimum for the nonbursty case, described in Section III. **B**. Optimal production rates at fixed *γ* = 1 for mean burst sizes *b* ∈ {0.5, 1, 5}, showing that increasing burst size lowers the overall rate magnitude. In both panels, mean FPT is fixed to *C* = 1 and the optimal rates are obtained numerically for the bursty model, explained in Section IV, without additional constraints on the production rates.

We finally studied the optimal production rate structure for higher threshold numbers. Fig. 7 shows the results for a ten-molecule threshold. Across the parameters considered, the optimal rates shape is nonmonotonic: production increases over intermediate molecule numbers before decreasing sharply near the threshold. Increasing the degradation rate raises the optimal production rates over much of the intermediate-state range while increasing the peak, consistent with the need to compensate for molecular loss (Fig. 7A). On the other hand, increasing the mean burst size lowers the overall magnitude of the optimal rates, which may be explained by the fact that each synthesis event produces more molecules on average and can advance the system across several states (Fig. 7B).

**Fig. 7:**
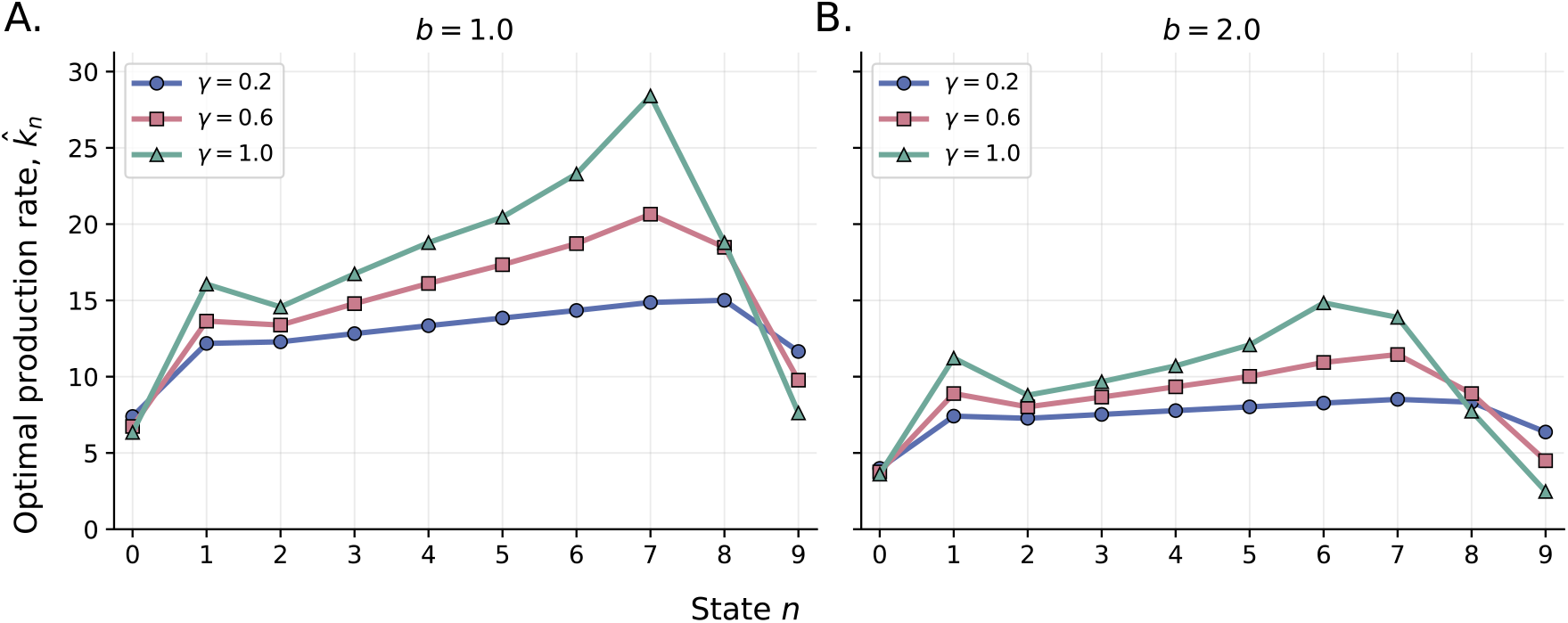
Degradation and burst size jointly produce a nonmonotonic optimal production rate structure for a ten-molecule threshold in bursty gene expression model. Optimal production rates are obtained numerically by solving the Lagrange derivative equations of the bursty model, explained in Section IV, subject to a fixed mean FPT *C* = 1 with no additional constraints imposed on the production rates. **A**. and **B**. show the optimal production rates for mean burst sizes *b* = 1 and *b* = 2, respectively, with degradation rates *γ* ∈ {0.2, 0.6, 1*}*.

The numerical optimization of the bursty model indicates that as the degradation rate is increased further, the largest optimal production rate grows rapidly and eventually becomes unbounded, indicating that a finite optimum is no longer obtained beyond a critical degradation rate. This behavior motivates examining the largest optimal rate, denoted by 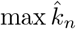 for 0 *≤ n ≤ X −* 1, directly as a function of *γ*. In Fig. 8, we extend the numerical continuation over a broader range of degradation rates to characterize the onset of this unbounded behavior and its dependence on the burst size and threshold. The results show that the largest optimal production rate increases with the degradation rate and rises sharply as a critical value is approached. Beyond this point, the numerical optimization no longer yields a finite maximum production rate. The transition occurs at lower degradation rates for larger thresholds, while its location is also altered by the mean burst size. Thus, the onset of the unbounded regime depends jointly on degradation rate, mean burst size, and threshold.

**Fig. 8:**
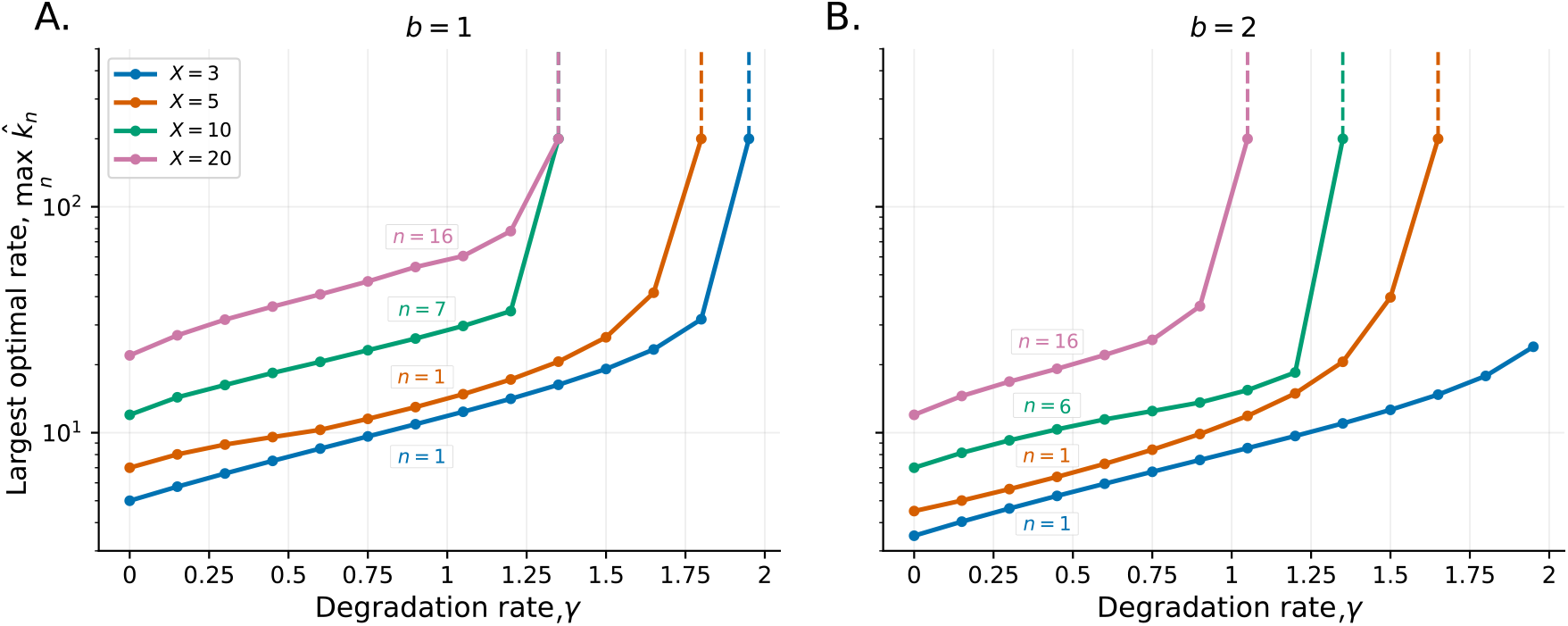
The largest optimal production rate increases with degradation and exhibits apparent divergence beyond a critical degradation rate. The largest optimal production rate, 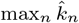 when 0 *≤n ≤X −*1, is shown as a function of the degradation rate *γ* for thresholds *X* ∈ {3, 5, 10, 20}. **A**. and **B**. correspond to mean burst sizes *b* = 1 and *b* = 2, respectively. The optimal rates are obtained numerically from the bursty fixed-mean FPT optimization problem, explained in Section IV, with *C* = 1. Solid curves show the finite optimal solutions. Dashed vertical lines mark the degradation rate beyond which the largest optimal production rate is no longer obtained as a finite numerical solution. The *n −* labels identify the state at which the largest optimal production rate is observed.

Overall, the bursty model retains a nonmonotonic optimal production rate structure across the finite-solution regime. Degradation rate and burst size mainly alter the magnitude and location of the peak, while sufficiently large degradation leads to the loss of a finite optimum. The onset of this unbounded regime depends also on both the molecule-number threshold and the mean burst size.

## V. Conclusions

In this work, we developed a framework for minimizing first-passage time variability in stochastic threshold-crossing systems with molecular degradation, considering both nonbursty and bursty gene expression. The problem was formulated using a birth-death model with arbitrary state-dependent production rates. Using backward equations and a Lagrange multiplier formulation, we derived optimality conditions for the production rates that minimize the FPT variability, subject to a fixed mean FPT.

The results show that degradation qualitatively reshapes the structure of the optimal feedback law. In the nonbursty production model, exact solutions were obtained for low thresholds, while asymptotic analysis for weak degradation at high thresholds showed that the optimal production rate increases approximately linearly with molecule number and drops sharply near the threshold. At larger degradation rates, the optimality conditions develop multiple solution branches and fold behavior, and the corresponding feedback structures can become oscillatory. The bursty production model further shows that randomness in production events can alter the optimal synthesis strategy, even at low thresholds, by changing how control effort is distributed across molecular states. At higher thresholds, the optimal production structures remain nonmonotonic, with degradation increasing the production rates required during the accumulation phase, while larger bursts reduce their overall magnitude. At sufficiently large degradation rates, however, the largest optimal production rate grows without bound, indicating the loss of a finite optimum. The onset of this unbounded regime depends on the molecule number threshold and mean burst size, as well as the degradation rate.

Together, these results demonstrate that degradation and burst statistics influence not only the magnitude of timing fluctuations and the shape of the optimal feedback structure, but also whether a finite optimal solution exists under a fixed-mean FPT constraint. The predicted state-dependent production structures could, in principle, be tested using synthetic gene circuits with tunable regulatory interactions, while single-cell measurements of threshold-crossing times could be used to compare the observed timing variability with the theoretical optimum. Future work could extend this framework to more mechanistic gene expression models, including additional sources of molecular noise and experimentally motivated regulatory architectures.

## Appendix A: Backward equations

We define the survival function [59] conditional on starting in state *n*. Survival is understood in the sense that the process has not crossed the threshold:

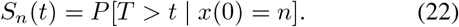

From first principles, survival over a short time interval is obtained by conditioning on the first transition: the process either jumps to *n−* 1 and survives, remains in *n* and survives, or jumps to *n* + 1 and survives. Thus,

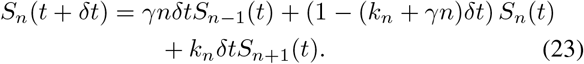

Passing to the limit *δt→* 0 in (23) yields the backward equation for the survival function,

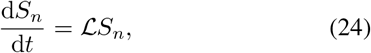

where *ℒ* is the generator of the Markov process *x*(*t*) [60]. This operator acts on sequences 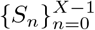 according to

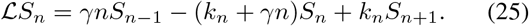

The boundary condition for (24) is *S*_*X*_ (*t*) = 0.

To derive equations for the FPT moments, consider the moment generating function (mgf) [61]

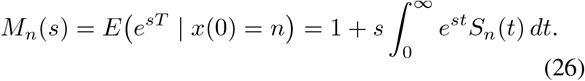

Multiplying (24) by *e*^*st*^ and integrating with respect to *t* yields the backward equation for the mgf,

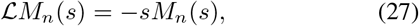

subject to the boundary condition *M*_*X*_ (*s*) = 1.

The FPT moments can be expressed in terms of the mgf as

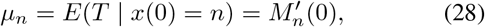

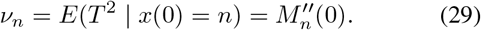

Differentiating (27) with respect to *s* and subsequently setting *s* = 0 yields

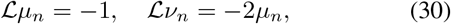

subject to boundary conditions *µ*_*X*_ = *ν*_*X*_ = 0. The first equation in (30), with ℒspecified by (25), coincides with equation (10) in the main text. After computing the mean FPTs *µ*_*n*_, these are substituted into the right-hand side of the equation for the second moments *ν*_*n*_.

## Appendix B: Regular perturbations

The Markov generator (25) can be expanded as

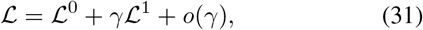

where

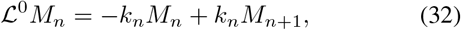

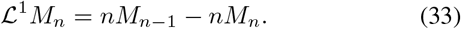

We expand the mgf as

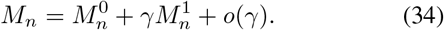

Substituting (31) and (34) into (27) and collecting terms of equal order yields

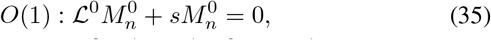

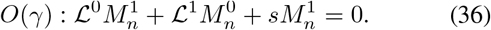

These equations are subject to the boundary condition 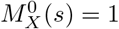 and 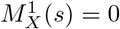.

Substituting (32) into (35)–(36) gives

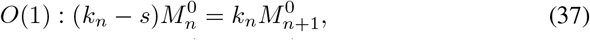

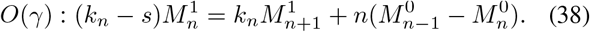

Equations (37)–(38) are first-order difference equations that can be solved by direct iteration in decreasing *n*. The zeroth component (corresponding to the zero initial state) is

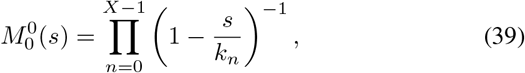

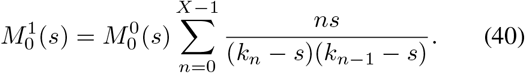

It is convenient to express the result in terms of the cumulant generating function (cgf) [61],

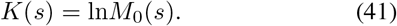

Substituting the perturbation expansion gives

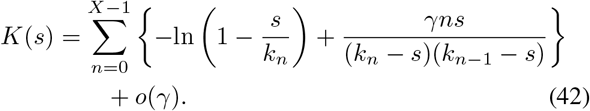

The mean and variance of the FPT can be expressed through the derivatives of the cgf [61]:

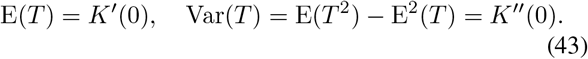

Differentiating (42) yields

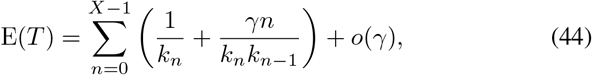

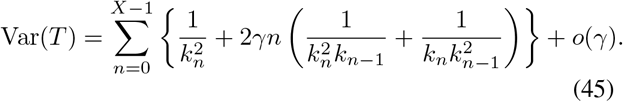

For analytical approximations, it is convenient to work with the Lagrangian equivalent to (6),

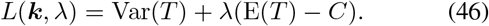

The corresponding Lagrange equations are

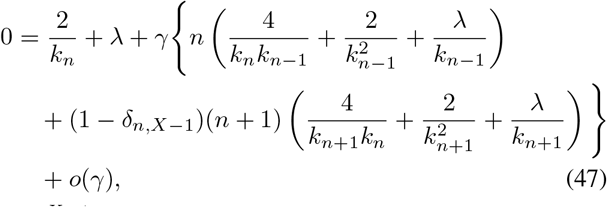

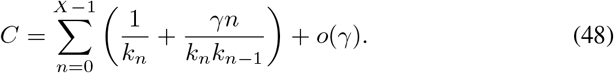

Solving (47)–(48) perturbatively yields the expressions (18)–(19).

## VI. Acknowledgments

This work is supported by NIH-NIGMS via grant R35GM148351.

## Notes

### Competing Interest Statement

The authors have declared no competing interest.

